# Optogenetic evaluation of the ability of different cutaneous C-fiber afferents to evoke aversive behaviors

**DOI:** 10.1101/2020.09.17.296384

**Authors:** Charles A. Warwick, Colleen Cassidy, Junichi Hachisuka, Margaret C. Wright, Kyle M. Baumbauer, Peter C. Adelman, Kuan H. Lee, Kelly M. Smith, Sarah E. Ross, H. Richard Koerber

**Affiliations:** Department of Neurobiology, University of Pittsburgh, 200 Lothrop St., Pittsburgh, PA 15213, USA; Pittsburgh Center for Pain Research, University of Pittsburgh, 200 Lothrop St., Pittsburgh, PA 15213, USA; Spinal Cord Group, Institute of Neuroscience and Psychology, University of Glasgow, Sir James Black Building, University Ave, Glasgow G12 8QQ, United Kingdom; Anatomy and Cell Biology, University of Kansas Medical Center, MS 3038, 3901 Rainbow Blvd., 2095 Hemenway, Kansas City, Kansas 66160

**Keywords:** Mrgd, SNI, electrophysiology, optogenetics, spared nerve injury, dorsal horn, spinoparabrachial, IB4

## Abstract

Most cutaneous C-fibers, including both peptidergic and non-peptidergic subtypes are presumed to be nociceptors and respond to noxious input in a graded manner. However, mechanically sensitive, non-peptidergic C-fibers also respond to mechanical input in the innocuous range, and so the degree to which they contribute to nociception remains unclear. To address this gap, we investigated the function of non-peptidergic afferents using the *Mrgprd^Cre^* allele. In real time place aversion studies, we found that low frequency optogenetic activation of *Mrgrpd^Cre^* lineage neurons was not aversive in naïve mice, but became aversive after spared nerve injury (SNI). To address the underlying mechanisms of this allodynia, we recorded from lamina I spinoparabrachial (SPB) neurons using the semi-intact *ex vivo* preparation. Following SNI, innocuous brushing of the skin gave rise to abnormal activity in lamina I SPB neurons, consisting of an increase in the proportion of recorded neurons that responded with excitatory post synaptic potentials or action potentials. This increase was likely due, at least in part, to an increase in the proportion of lamina I (LI) SPB neurons that received input upon optogenetic activation of *Mrgprd^Cre^* lineage neurons. Intriguingly, in SPB neurons there was a significant increase in the EPSC latency from *Mrgprd^Cre^* lineage input following SNI, consistent with the possibility that the greater activation post SNI could be due to the recruitment of a new polysynaptic circuit. Together, our findings suggest *Mrgprd^Cre^* lineage neurons can provide mechanical input to the dorsal horn that is non-noxious before injury but becomes noxious afterwards due the engagement of a previously silent polysynaptic circuit in the dorsal horn.

## INTRODUCTION

Traditionally, cutaneous C-fibers have been categorized into two major classes: peptidergic afferents that generally express substance P and/or CGRP, as well as the so-called non-peptidergic afferents that express Mrgprd and/or bind the isolectin IB4 [3; 9]. Because both populations respond vigorously to noxious stimuli, it was generally assumed that both types function mainly as nociceptors. Consistent with this idea, ablation studies have suggested that peptidergic afferents are responsible for thermal pain, whereas non-peptidergic afferents are responsible for mechanical pain [6]. Moreover, because these nociceptive-responsive neurons collectively make up the vast majority of C-fibers, it was not uncommon to generally equate C-fiber activity with nociception and, by extension, pain.

But other findings have challenged the presumption that these populations are exclusively pain-inducing nociceptors. In particular, it remains unclear whether the non-peptidergic subset of IB4-binding afferents signal nociception, or whether their function is more nuanced. For instance, although non-peptidergic, IB4-binding neurons respond to noxious stimuli, they typically show a graded response over a wide range of mechanical forces, and indeed have the capacity to detect even low threshold input that is clearly non-noxious [21]. In support of this idea, human microneurography studies have shown that the vast majority of mechanically sensitive C-fibers are activated by stimulus intensities that are reported as non-painful [12; 27]. Finally, although transient optogenetic activation of Mrgprd neurons in mice was found to cause withdrawal, prolonged exposure was not sufficient to elicit conditioned place aversion [4]. Thus, the degree to which activity in the non-peptidergic, IB4-binding population is sufficient to drive aversion and/or pain remains ambiguous.

A second major gap in our understanding is the role of non-peptidergic afferents in chronic pain. Injuries that give rise to chronic pain, such as spared nerve injury (SNI), cause central changes that alter the way sensory information is integrated by the nervous system [2; 28]. One of the most common consequences of these injury-induced changes is allodynia: a phenomenon in which innocuous mechanical stimuli, such as light brushing of the skin, are perceived as noxious. It is generally assumed that the afferents responsible for allodynia are those that are most responsive to innocuous mechanical input, *i.e.* low threshold mechanoreceptors (LTMRs). However, many non-peptidergic IB4-binding neurons frequently respond vigorously to innocuous stimuli, such as brushing, raising the possibility of their involvement in allodynia [21]. Thus, there remains significant uncertainty about both the afferent subtype(s) and the spinal circuits downstream thereof that give rise to allodynia.

Here, we used the *Mrgprd^Cre^* allele to investigate the function of non-peptidergic IB4-binding C-fibers in normal sensation and in chronic neuropathic pain. In a real time place aversion assay, we show that activation of *Mrgprd^Cre^* lineage neurons is not aversive in naïve mice. However, in the context of SNI-induced neuropathic pain, activation of *Mrgprd^Cre^* lineage neurons became aversive. We subsequently show that SNI increases the proportion of lamina I SPB neurons that respond to innocuous brush input, and that this increase is likely to be mediated, at least in part, by the activity of *Mrgprd^Cre^* lineage neurons, which appear to gain access to lamina I SPB neurons by way of an emergent polysynaptic circuit following SNI.

## METHODS

*Animals* The mice used in these experiments were *Mrgprd^tm1.1(cre)An^* [21], which were obtained from the MMRRC at Chapel Hill (Stock No: 036118-UNC); *Trpv1^tm1(cre)Bbm^* [5], which were obtained from Jax labs (Stock No: 017769) and *Gt(ROSA)26Sor^tm32(CAG-CAP4*H134R/EYFP)Hze^* also known as Ai32 mice [17], which were obtained from Jax labs (Stock No: 024109). Additional crosses were made to obtain *Mrgprd^Cre^;* Ai32 and *Trpv1^Cre^;* Ai32 mice for these experiments. Mice were given free access to food and water and housed under standard laboratory conditions. The use of animals was approved by the Institutional Animal Care and Use Committee of the University of Pittsburgh.

### Ex vivo preparation

The *ex vivo* somatosensory system preparation has been described previously in detail [10; 18]. Briefly, adult mice (>4 weeks of age) were anesthetized with a mixture of ketamine and xylazine (90 and 10 mg/kg, respectively) and perfused transcardially with room-temperature, oxygenated (95% O_2_–5% CO_2_) artificial CSF (aCSF) (in mmol/L: 1.9 KCl, 1.2 KH_2_PO_4_, 1.3 MgSO_4_, 2.4 CaCl_2_, 26.0 NaHCO_3_, and 10.0 D-glucose), with 253.9 mmol/L sucrose. Spinal cord, L1–L4 dorsal root ganglia (DRG), saphenous nerve, and innervated skin were dissected free in continuity. After dissection, the preparation was transferred to a separate recording chamber containing oxygenated aCSF in which the sucrose was replaced with 127.0 mmol/L NaCl. The skin was pinned out on a stainless-steel grid platform located at the bath/air interface, such that the dermal surface remained perfused with the aCSF while the epidermis was exposed to the air. The platform provided stability during application of thermal and mechanical stimuli. The bath was then slowly warmed to 31 °C before recording.

### DRG recordings and peripheral stimuli

Intracellular recordings from L3 DRG cells in the were made using quartz microelectrodes, (> 100 MΩ filled with 1 mol/L K-acetate). An electric search stimulus was applied at 1.5 Hz through a glass suction electrode applied to the saphenous nerve to locate cells with axons in the saphenous nerve. Cutaneous receptive fields (RFs) were located with a fine paint brush, blunt glass probe, and von Frey hairs. When cells were driven by the nerve but had no mechanical RF, a thermal search was performed by gently applying hot (52°C) and cold (0°C) saline to the surface of skin using a syringe with a 20-gauge needle. If a thermal RF was located, the absence of mechanical sensitivity was confirmed by searching the identified RF using a glass probe. In most cases, the response characteristics of the DRG cell were then determined by applying computer-controlled mechanical and thermal stimuli (Fig. 2A). The mechanical stimulator consisted of a constant-force controller (Aurora Scientific) attached to a 1-mm-diameter plastic disc. Computer-controlled 5 s square waves of 5, 10, 25, 50, and 100 mN were applied to the RF of the cell. Following the functional characterization, responsiveness to laser stimulation was determined using an 80-mW, 473-nm wavelength laser and a 200-micron fiber optic cable affixed to a micromanipulator (Laserglow Technologies, Toronto, Canada). The opening of the fiber optic cable was positioned approximately 5 mm from the skin surface.

### Parabrachial injections

Four- to 6-week-old mice were anesthetized with isoflurane and placed in a stereotaxic apparatus. An incision was made to expose the bone and a small hole was made in the skull with a dental drill. A glass pipette was used to inject 100 nL of FAST DiI oil (2.5 mg/mL; Invitrogen, Carlsbad, CA) into the left lateral parabrachial area at the following coordinates: from lambda, 1.3 mm lateral; from lambdoid suture, 0.5 mm posterior; from surface, −2.4 mm. The incision was closed with sutures and the mice were allowed to recover and returned to their home cages. These injections were made at least 5 days before electrophysiological recordings.

### Whole-cell spinal neuron recordings

Neurons were visualized using a fixed-stage upright Olympus microscope equipped with a 40x water immersion objective, a CCD camera (ORCA-ER; Hamamatsu Photonics, Hamamatsu City, Japan) and monitor. A narrow-beam infrared LED (L850D-06; Marubeni, Tokyo, Japan, emission peak, 850 nm) was positioned outside the bath, as previously described [10]. Projection neurons in lamina I were identified by DiI fluorescence following retrograde labeling in the parabrachial nucleus. Whole-cell patch clamp recordings were made using borosilicate glass microelectrodes pulled using a PC-10 puller (Narishige International, East Meadow, NY). Pipette resistances ranged from 6 to 12 MΩ. Electrodes were filled with a solution containing the following (in mM): 135 K-gluconate, 5 KCl, 0.5 CaCl2, 5 EGTA, 5 HEPES, and 5 MgATP; pH 7.2. Alexa fluor 488 (Invitrogen; 25 mM) was added to confirm recording from the targeted cell. Recordings were acquired using an Axopatch 200B amplifier (Molecular Devices, Sunnyvale, CA). The data were low-pass filtered at 2 kHz and digitized at 10 kHz using a Digidata 1322A (Molecular Devices) and stored using Clampex version10, (Molecular Devices).

### Cutaneous stimulation

The cell’s receptive field was determined using a paint brush and 1- to 4-g von Frey filaments. Once the cell’s receptive field was located, subsequent stimuli were applied directly to the receptive field for 1 s to determine the cell’s response properties. MiniAnalysis (Synaptosoft, Decatur, GA) was used for detecting excitatory post synaptic currents (EPSCs) and action potentials (APs).

### Cell dissociation and pickup for RT-PCR

DRG containing labeled cells from *Mrgprd^Cre^;* Ai32 mice were removed and dissociated as described previously [1]. Briefly, DRG were treated with papain (30 U) followed by collagenase CLS2 (10 U)/Dispase type II (7 U), centrifuged (1 min at 1000 r/min), triturated in Minimal Essential Medium, plated onto laminin-coated coverslips in 30 mm diameter dishes, and incubated at 37 °C for 45 min. Dishes were removed and flooded with collection buffer (140 mM NaCl, 10 mM Glucose, 10 mM HEPES, 5 mM KCl, 2 mM CaCl_2_, 1 mM MgCl_2_). Single, labeled cells were identified using fluorescence microscopy, picked up using glass capillaries (World Precision Instruments) held by a 4-axis micromanipulator under bright-field optics, and transferred to tubes containing 3 μL of lysis buffer (Epicentre, MessageBOOSTER kit). Cells were collected within 1 h of removal from the incubator and within 4 h of removal from the animals.

### Single-cell amplification and qPCR

The RNA isolated from each cell was reverse transcribed and amplified using T7 linear amplification (Epicentre, Message BOOSTER kit for cell lysate), run through RNA Cleaner & Concentrator-5 columns (Zymo Research), and analyzed using qPCR, as described previously [1], using optimized primers and SsoAdvanced SYBR Green Master Mix (Bio-Rad). Threshold cycle time (Ct) values were determined for each well.

### Real time place aversion (RTPA)

Behavioral boxes were constructed from plexiglass that were 12 (l) x 5 (w) x 10 (h) inches with guides for dividing the boxes into three chambers: 2 (5 x 5 inches) and 1 (2 x 5 inches) in the middle. For optical stimulation, we used Blue LED Strip Light, 120/m, 10 mm wide, by the 5m Reel, 12 VDC, 8.9 Watts/meter, 742 mA/meter (centered at 460 nM) and Amber LED Strip Light, 120/m, 10 mm wide, by the 5m Reel, 12 VDC, 8.9 Watts/meter, 742 mA/meter (centered at 595 nM) (Environmental Lights, San Diego, California). Five-inch strips of LEDs were affixed to a 5- X 12-inch metal box (heat sink) covering a 5- x 5-inch area on the ends of the box (one area blue the other amber) leaving a 2-inch area in the middle free of lights. This box could then be placed under the plexiglass box to illuminate the floor (Fig. 3D). The LEDs were controlled using a CED 1401 interface. The LEDs were positioned 10 mm below the surface of the plexiglass floor as we found that at this distance there was no heating of the plexiglass floor over the 15-min period of observation.

Mice used in the RTPA were first acclimated to the behavioral boxes for 3 days (15 minutes/day) before testing. On the day of testing the mice were placed in the center compartment and the LEDs turned on and the sliding doors removed so the mice could move freely throughout the box for 15 min. The mice were recorded during this period using a video camera. The videos were then scored off line. All experimenters were blinded to genotype and experimental condition of the mice.

### Spared nerve injury

Mice were anesthetized with isoflurane and the posterior right hindlimb was shaved and cleaned using an aseptic solution. An incision was made over the popliteal fossa and the underlying muscle incised to expose the fossa. The tibial and peroneal nerves were isolated. The two nerves were tightly ligated and cut just distal to the ligature. The muscle and skin were sutured and the animal allowed to recover. For sham surgeries the procedures were the same except for the ligation and transection.

### Immunohistochemistry

Deeply anesthetized mice were perfused with 4% paraformaldehyde and tissue sections of the L3 spinal cord, DRG, and glabrous skin were embedded in OCT and cut on a cryostat at 20 μm. The sections were stained for IB4, CGRP and eYFP using the following antibodies or probes: anti-GFP (chicken 1:1250; Aves Labs), anti-CGRP (rabbit; 1:1000; Chemicon) and IB4 (1:250; IB4-congugated AlexaFluor 647; Molecular Probes, Eugene, OR). After incubation in primary antiserum for at least 2 h, tissue was washed and incubated in appropriate fluorescently tagged secondary antibodies (1:500; Jackson Immunoresearch) for 1 h followed by Hoescht staining (1:10,000) or 30 min. After washing and mounting in Fluoromount (Sigma), sections were viewed and imaged on an Olympus BX53 fluorescent microscope with UPanSApo 10x or 20x objectives.

### Statistical analyses

Data are expressed as the mean ± SEM. For electrophysiological experiments, dots represent data points from individual neurons combined from all animals. The following tests were used for statistical analysis: unpaired Student’s *t* for continuous normal data (comparison of 2 groups), Chi-squared or Fisher’s exact test for categorical data, ordinary one-way ANOVA with Tukey’s multiple comparisons *post hoc* test if indicated by the main effect (comparison of >2 groups), or two-way repeated-measures ANOVA with Tukey’s multiple comparisons *post hoc* test if indicated by the main effect (time course with comparison of 2 or more groups). All tests were two-tailed (unless otherwise indicated) and a value of P < 0.05 was considered statistically significant in all cases. All statistical analyses were performed using GraphPad Prism 8.4 software.

## RESULTS

### Histological and molecular characterization of Mrgprd^Cre^ lineage neurons

To investigate the functional role IB4-binding neurons, we targeted these neurons genetically using the *Mrgprd^Cre^* knockin allele that was originally developed by David Anderson [21]. Because Mrgprd shows some developmental expression [16], we carefully characterized the neurons that are captured by this genetic tool, as assessed using Ai32, a Cre-dependent reporter embedded in the *Rosa* locus that enables expression of a ChR2-eYPF fusion protein [17]. Analysis of cell bodies in lumbar DRG suggested that the *Mrgprd^Cre^* allele captured ~80% of IB4-binding neurons, and similarly that ~80% of IB4-binding neurons expressed ChR2-eYFP (Figs. 1A-C). In contrast, the majority of the CGRP-expressing afferents did not express ChR2-eYPF (Figs. 1D-E). The analysis of the innervation pattern in the lumbar spinal cord showed a similar pattern: *Mrgprd^Cre^* lineage neurons showed a very high degree of colocalization with IB4 in a band that corresponds approximately to inner lamina II, but limited colocalization with CGRP, which primarily targets lamina I and outer lamina II (Figs. 1F-J). In the glabrous skin, *Mrgprd^Cre^*-targeted neurons colocalized with the pan-neuronal marker PGP9.5, and innervated the most superficial aspect of the epidermis (Figs. 1K-M).

**Figure 1.**
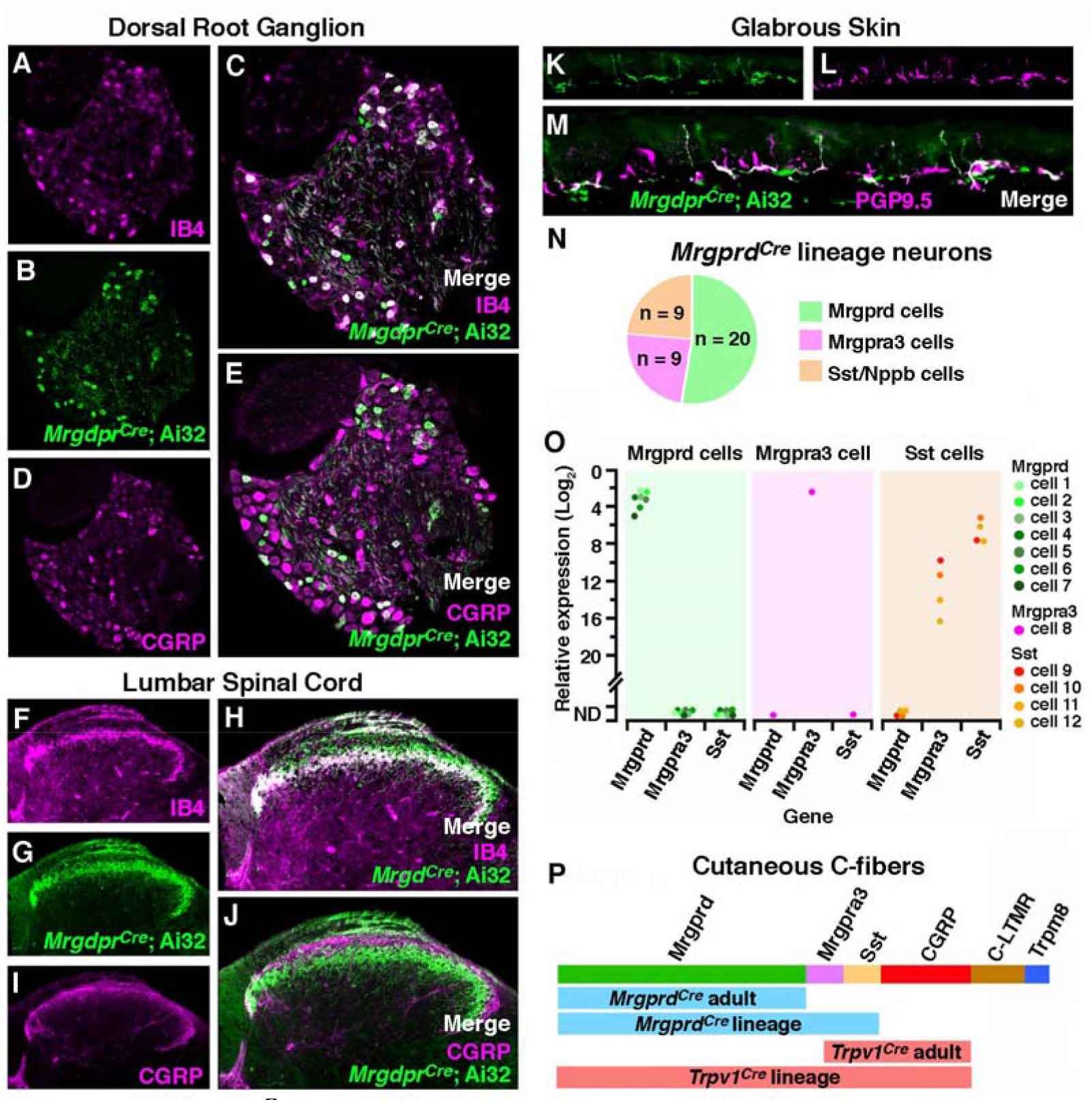
*Mrgprd^Cre^ lineage afferents, which comprise Mrgprd, Mrgpra3, and Sst populations, target lamina IIi in the dorsal horn and the superficial aspect of the skin.* (A - E) Representative images of a lumbar dorsal root ganglion from a *Mrgprd^Cre^;* Ai32 mouse that is co-stained for eYFP and IB4 or CGRP. (F – J) Representative images of the superficial dorsal horn, lumbar level from a *Mrgprd^Cre^;* Ai32 mouse that is co-stained for eYFP and IB4 or CGRP. (K – M) Representative images of the glabrous skin from a *Mrgprd^Cre^;* Ai32 mouse that is co-stained for eYFP and PGP9.5. (N) Summary of the classification of *Mrgprd^Cre^* lineage neurons based on single cell RT-PCR (O) Expression of *Mrgprd, Mrgpra3, and Sst* mRNA relative to *Gapdh* in individual eYFP-marked *Mrgprd^Cre^* lineage neurons. Data are presented as the log_2_ ΔCT expression relative to *Gapdh* expression within the same cell such that smaller numbers represent higher mRNA expression. Colored dots represent data points from individual cells from a single, representative mouse. (P) Schematic to illustrate the different subtypes of cutaneous C-fibers that are captured by the *Mrgprd^Cre^* and *Trpv1^Cre^* alleles with developmental (lineage) or adult recombination.

A number of recent single-cell sequencing studies have suggested that non-peptidergic afferents can be further subdivided into more refined subtypes characterized by expression of *Mrgprd, Mrgpra3,* or *Sst/Nppb* [20; 23; 26]. To assess whether the *Mrgprd^Cre^* lineage captures these populations, we performed quantitative real-time PCR of individual eYFP-labeled DRG neurons that were freshly dissociated from adult *Mrgprd^Cre^;* Ai32 mice. These experiments revealed that all genetically-marked neurons fell into one of three apparent cell types: those with high *Mrgprd* (and lacking *Mrgpra3, Sst, Calca* and *Tac1);* those with high levels of *Mrgpra3* (as well as *Calca,* but little or no *Mrgprd, Sst* and *Tac1*); and those with high levels of *Sst* (and low/moderate levels of *Mrgpra3* and *Tac1,* but lacking *Mrgprd, Calca)* (Fig. 1N). Importantly, this single-cell analysis of eYPF-labeled cells indicated that *Mrgprd* is no longer detected in the *Mrgpra3* or *Sst* populations in the adult, suggestive of developmental expression (Fig. 1O). Thus, the *Mrgprd^Cre^* lineage faithfully captures the majority of afferents that have traditionally been termed ‘non-peptidergic’ nociceptors, which includes most IB4-binding neurons, but not peptidergic nociceptors, C-LTMRs, or Trpm8-expressing cold/cool-detectors. For purposes of comparison, we also visualized the recombination mediated by the *Trpv1^Cre^* allele. Consistent with previous reports [5], we observed that *Trpv1^Cre^* lineage neurons include the vast majority of cutaneous C-fibers, though not C-LTMRs or Trpm8-expressing afferents (summarized in Fig. 1P).

### *Optogenetic manipulation of Mrgprd^Cre^* lineage neurons

We next characterized the responses of *Mrgprd^Cre^* lineage neurons to optogenetic stimulation of the skin with *Mrgprd^Cre^;* Ai32 mice and the *ex vivo* preparation, performing intracellular recordings of L3 DRG neurons (Fig. 2A). Afferents were categorized as cutaneous C-fibers if they responded to electrical stimulation of the saphenous nerve with a conduction velocity of 1.2 m/s or less. Next, the receptive field of a given cutaneous C-fiber was identified manually through application of mechanical and/or thermal stimuli. Thereafter, quantitative phenotyping to natural and optogenetic stimuli was performed using a Peltier thermode, a mechanical stimulator, and a 473-nm laser applied to the skin within the identified receptive field. Typically, *Mrgprd^Cre^* lineage neurons responded weakly to cooling and vigorously to noxious heat; they also responded to both low threshold (10 mN) and high threshold (50 mN) mechanical input in a graded manner, consistent with previous characterizations of non-peptidergic nociceptors [14; 21]. Both optogenetic and natural stimulation of the skin elicited action potentials (Fig. 2D). We found that a 5-ms pulse of light was sufficient to drive a single action potential, whereas a 300-ms pulse of light was required to observe a doublet. Finally, *Mrgprd^Cre^* lineage neurons could follow optogenetic stimulation of at least 5 Hz (Fig. 2C).

**Figure 2.**
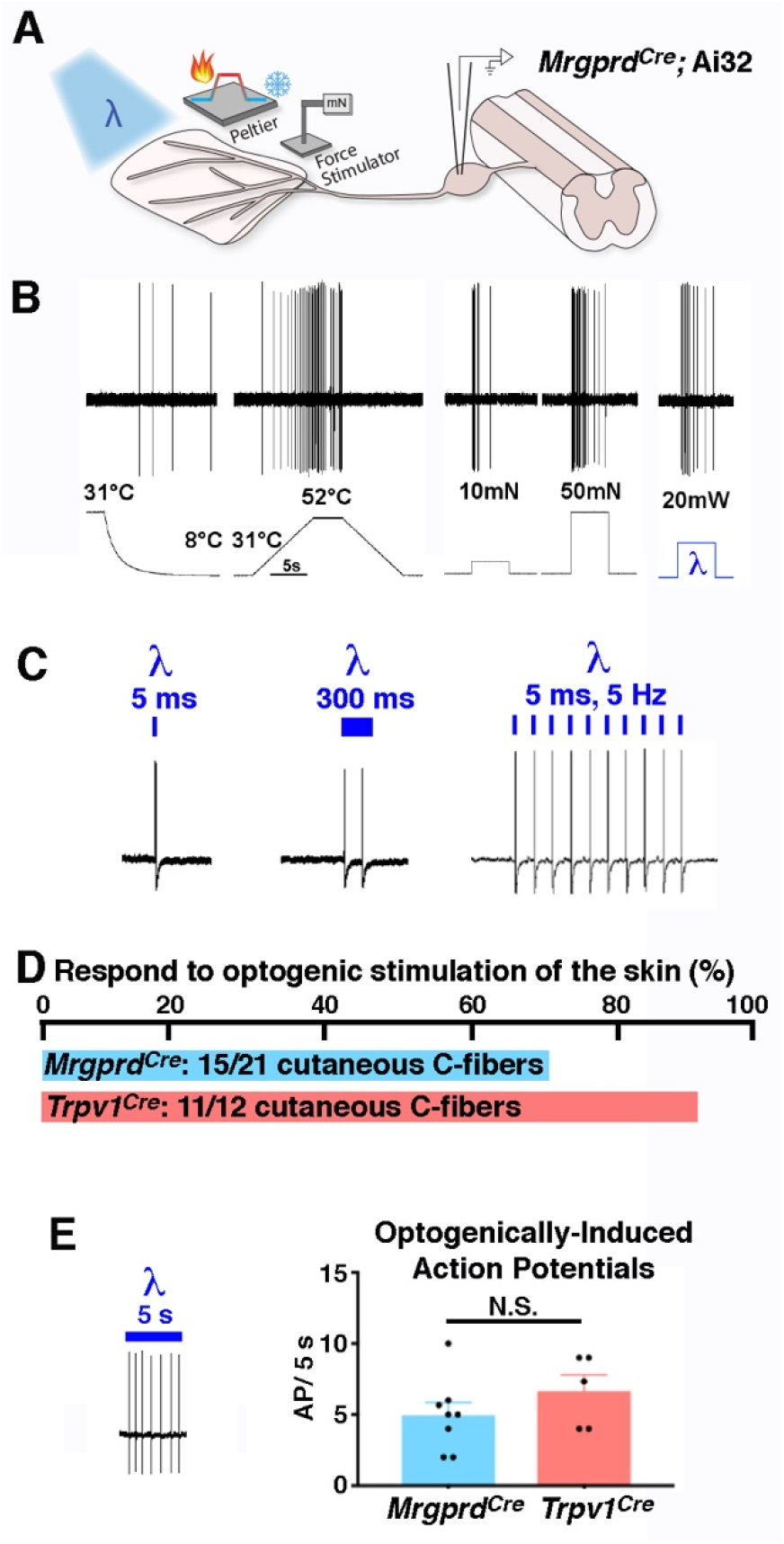
*Response of Mrgprd^Cre^ lineage afferents to optogenetic stimulation of the skin.* (A) Schematic of cutaneous stimulation and intracellular DRG recordings using the *ex vivo* preparation. (B) Representative responses of *Mrgprd^Cre^* lineage afferents to cooling, heating, mechanical and optogenetic stimulation. (C) Representative responses to optogenetic stimulation of *Mrgprd^Cre^* lineage afferents of varying duration and frequency. (D) Proportion of C-fibers that respond to optogenetic stimulation of the skin in *Mrgprd^Cre^;* Ai32 and *Trpv1^Cre^;* Ai32 mice, as indicated. (E) Representative recording and quantification of the number of action potentials observed in a given cell upon optogenetic stimulation of the skin in *Mrgprd^Cre^;* Ai32 and *Trpv1^Cre^;* Ai32 mice, as indicated. Data are mean ± SEM with dots representing data from individual neurons; N.S. indicates p > 0.05 (unpaired Student’s t-test).

Overall, of 21 cutaneous C-fibers that were recorded from *Mrgprd^Cre^;* Ai32 mice, 15 showed responses to optogenetic stimulation (~70%), consistent with the idea that *Mrgprd^Cre^* lineage neurons represent the majority of cutaneous C-fibers. For comparison, we found that 11 of 12 (~90%) cutaneous C-fibers from *Trpv1^Cre^;* Ai32 mice could be opto-tagged in this fashion (Fig. 1D), as expected given the broad recombination mediated by this allele [5]. Importantly, though the *Trpv1^Cre^* allele captured slightly more C-fibers than the *Mrgprd^Cre^* allele, optogenetic stimulation of individual C-fibers from mice of either genotype gave rise to similar numbers of action potentials, thereby enabling direct comparison of the two alleles in behavioral studies (Fig. 1E).

### *Behavioral responses to optogenetic manipulation of Mrgprd^Cre^* lineage neurons

Withdrawal in animals has traditionally been interpreted to represent a pain behavior, but the percept associated with this response is difficult to infer. Previous work has shown that selective optogenetic activation of Mrgprd neurons elicits withdrawal, but not flinching, licking, or guarding [4]. We anticipated that optogenetic stimulation of *Mrgprd^Cre^* lineage neurons would be at least as aversive, if not more so, because the genetic tool we used captures a significantly larger number of C-fibers than that used by Beaudry et al. [4]. Although both alleles target the same genetic locus, the degree of recombination using the *Mrgprd^Cre^* allele is more extensive than the *Mrgprd^CreER^* allele for two reasons: the constitutive Cre allele captures more than the adult Mrgprd population due to developmental expression (i.e., Sst and Mrgpra3 populations), whereas the *Mrgprd^CreER^* allele likely captures less than the adult Mrgprd population due to incomplete recombination. Contrary to our expectations, however, we found that the responses of *Mrgprd^Cre^* mice to optogenetic stimulation of the hindpaws appeared quite modest, and not qualitatively different than that described for the *Mrgprd^CreER^* mice [4]. Resting mice generally withdrew their hindpaw for a brief moment, as if startled, and then replaced it on the floor, and alert mice rarely responded at all. In sharp contrast, activation of *Trpv1^Cre^* lineage neurons invariably caused rapid, robust withdrawal that was frequently accompanied by flinching and/or licking (Fig. 3A), just as previously described [4]. These observations raised the possibility that low-frequency optogenetic activation of non-peptidergic ‘nociceptors’ might not actually be aversive.

**Figure 3.**
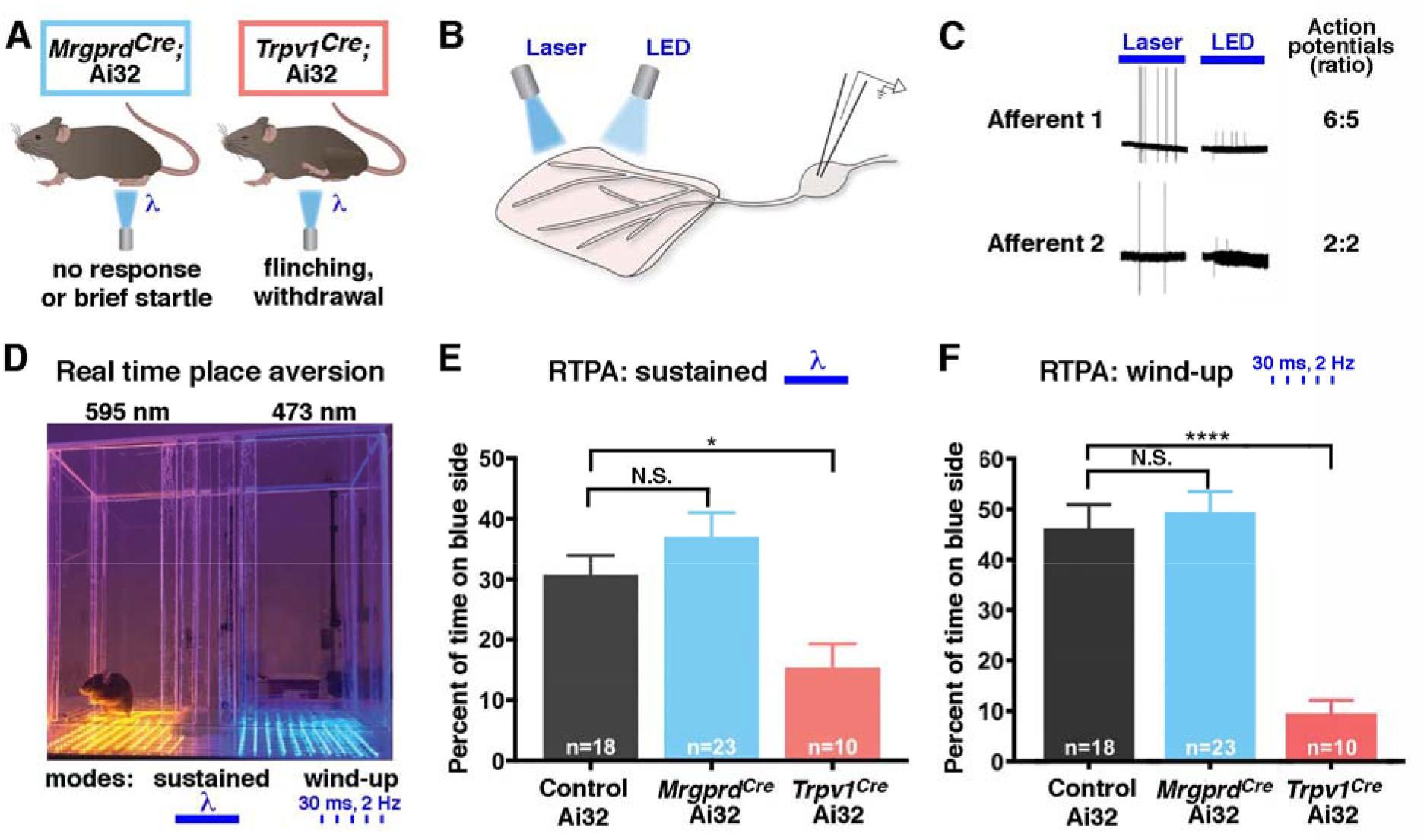
*Low-frequency, cutaneous optogenetic activation is aversive in Trpv1^Cre^ mice but not Mrgprd^Cre^ mice in the naïve condition.* (A) Schematic illustrating behavioral responses to optogenetic stimulation at the hindpaw of *Mrgprd^Cre^;* Ai32 and *Trpv1^Cre^;* Ai32 mice. (B) Schematic illustrating comparison of laser- and LED-mediated optogenetic stimulation. (C) Representative traces and quantification of laser-evoked and LED-evoked action potentials in *ex vivo* recordings from *Mrgprd^Cre^;* Ai32 mice. (D) Picture of optogenetic real time place aversion (RTPA) apparatus with amber-light and blue-light sides separated by a middle chamber. (E) In response to sustained light, *Trpv1^Cre^;* Ai32 mice, but not *Mrgprd^Cre^;* Ai32 mice, show real time place aversion to the blue-light side of the apparatus. Data are mean ± SEM; n is number of mice; N.S. indicates P > 0.05; * indicates the multiple comparison adjusted P-value < 0.05 relative to Control Ai32. Significant main effect: F = 6.793, P < 0.0025 (Ordinary one-way ANOVA with Tukey’s multiple comparisons post hoc test). (F) In response to flashing light (2 Hz, wind-up mode), *Trpv1^Cre^;* Ai32 mice, but not *Mrgprd^Cre^;* Ai32 mice, show real time place aversion to the blue-light side of the apparatus. Data are mean ± SEM; n is number of mice; N.S. indicates multiple comparison adjusted P-value > 0.05; **** indicates multiple comparison adjusted p-value < 0.001 relative to Control Ai32. Significant main effect: F = 18.28, P < 0.0001 (Ordinary one-way ANOVA with Tukey’s multiple comparisons post hoc test).

Because reflexive behaviors can be hard to interpret and do not readily assess aversion, we turned to non-reflexive behaviors for quantitative analysis. When a stimulus is quite noxious, such as intraplantar formalin, mice will develop conditioned place aversion to the side of the chamber that is associated with the aversive experience. Intriguingly, Beaudry et al. showed that optogenetic activation of *Trpv1^Cre^* afferents was sufficient for conditioned place aversion, whereas activation of *Mrgprd^CreER^* afferents was not [4]. However, conditioned place aversion can be difficult to achieve unless the paired stimulus is fairly injurious. We therefore developed a real time place aversion (RTPA) assay to enable the detection of aversive stimuli with higher sensitivity.

Our previous optogenetic characterization had been done by laser, a coherent light source that is ideal for focal stimulation. However, for the RTPA studies we were planning to use LEDs, an incoherent light source which is more suitable for widespread activation. To compare the efficacy of optogenetic stimulation by laser and LED, we performed *ex vivo* skin nerve recordings (Fig. 3B). Importantly, stimulation of the skin with either light source gave rise to a similar number of action potentials in ChR2-expressing *Mrgprd^Cre^* lineage neurons, indicating that optogenetic stimulation by laser or LED was equally effective (Fig. 3C). With this knowledge, we built a custom RTPA box comprising a stimulation side in which the floor was lined an array of blue LED lights and a control side in which the floor was lined an array of amber LED lights, which were separated by a small, unlit middle chamber. These lights were set to run in either the sustained mode, in which the lights were constantly on, or the ‘wind-up’ mode, in which the lights would flash at 2 Hz for 30 ms per flash. Mice were placed in the RTPA box for 15 minutes, during which time they were allowed to move freely from one chamber to another (Fig. 3D).

Next, we performed RTPA assays to quantify the aversiveness of optogenetic stimulation. Just as expected, we found that *Trpv1^Cre^;* Ai32 mice showed significant avoidance of the blue-light side, consistent with the idea that optogenetic activation of nociceptors is unpleasant. In sharp contrast, the time that *Mrgprd^Cre^;* Ai32 mice spent on the blue-light side was not different than control littermates harboring Ai32 alone (Fig. 3E). Similar findings were observed when the LED lights were flashing at 2 Hz to simulate wind-up (Fig. 3F). These striking observations suggest that optogenetic activation of *Mrgprd^Cre^;Ai32* fibers was not aversive, at least not in naïve mice.

### Effect of injury: behavioral responses to optogenetic manipulation of Mrgprd^Cre^ lineage neurons in the context of neuropathic pain

Although our data suggested that *Mrgprd^Cre^* lineage neurons can be activated without causing acute pain, a key remaining question was whether they contribute to chronic pain. Allodynia is one of the hallmarks of chronic pain, particularly neuropathic pain [13]. For people that suffer from allodynia, innocuous everyday experiences, such as fabric moving across the skin, can give rise to shooting pain. Since many non-peptidergic afferents can respond robustly to dynamic low threshold stimulation [21], we hypothesized that *Mrgprd^Cre^* lineage neurons might contribute to this allodynia. To address this question, we selected to use the spared nerve injury (SNI) model [8] of chronic neuropathic pain (Fig. 4A). Although the saphenous nerve remains intact in this model, there is nevertheless the potential for the function of afferents to be affected by the injury. We therefore measured the responses of *Mrgprd^Cre^* lineage afferents to optogenetic stimulation of the skin, comparing naïve and SNI mice (Fig. 4B). Importantly, the number of action potentials that were elicited was not affected by SNI (Fig. 4C). Thus, neuropathic injury did not cause sensitization of *Mrgprd^Cre^* lineage afferents to optogenetic stimulation.

**Figure 4.**
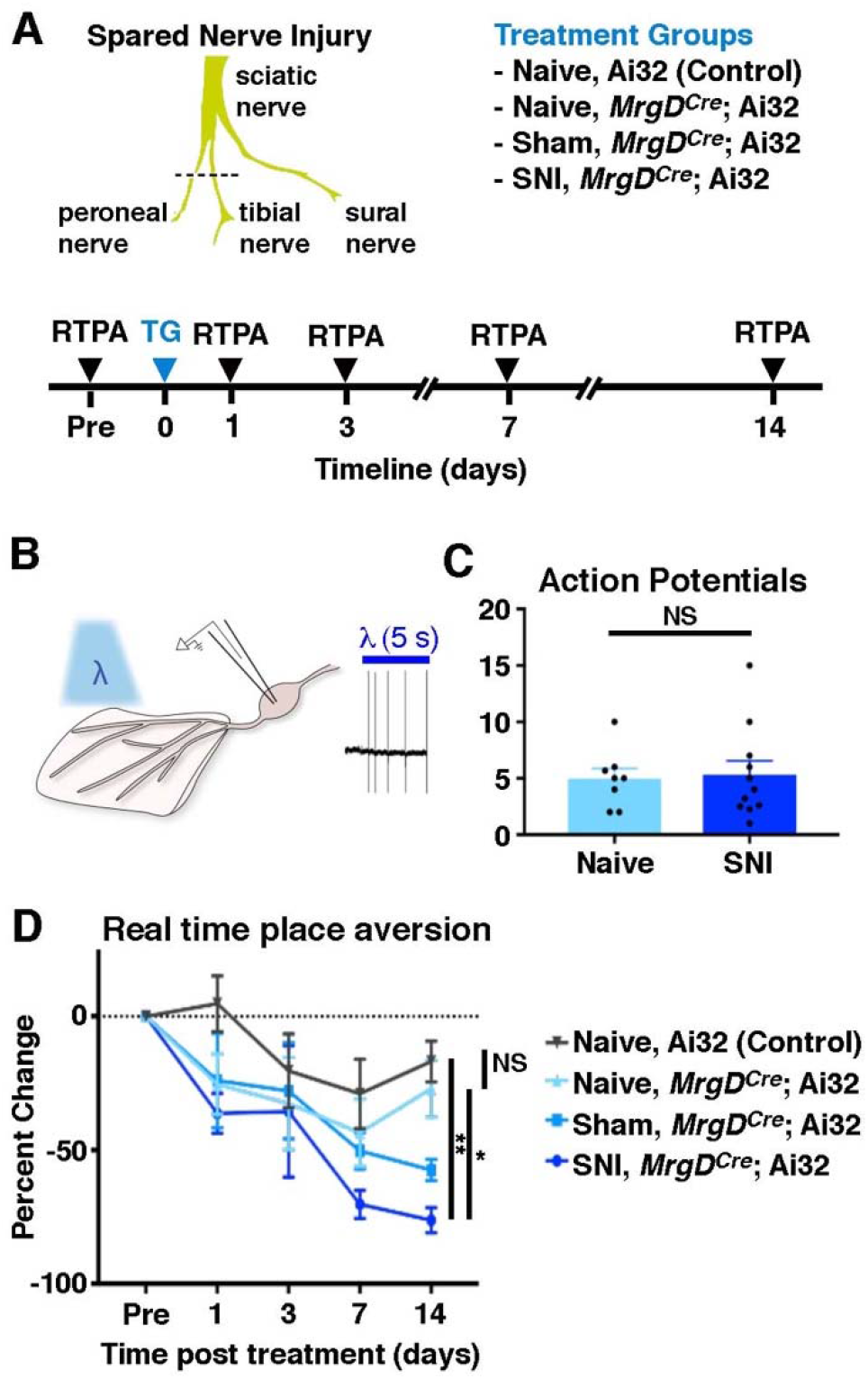
Optogenetic activation of *Mrgprd^Cre^* lineage afferents is aversive after SNI. (A) Schematic illustrating experimental design. (B) Schematic and representative recording of responses of opto-tagged C-fibers to cutaneous optogenetic stimulation in naïve and SNI mice. (C) Quantification of the number of action potentials observed upon optogenetic stimulation of the skin in individual neurons from naïve and SNI *Mrgprd^Cre^*; Ai32 mice. Data are mean ± SEM with dots representing data from individual neurons; N.S. indicates P > 0.05 (unpaired Student’s t-test). (D) Real time place aversion across four treatment groups: naïve controls (no ChR2); naïve *Mrgprd^Cre^;* Ai32; sham *Mrgprd^Cre^;* Ai32; and SNI *Mrgprd^Cre^;* Ai32. Data are mean ± SEM. n = 5 - 6 mice per group. There was a significant main effect of Time: F (4, 76) = 14.82, P < 0.0001; Treatment groups: F (3, 19) = 3.566. P = 0.0336; and Subject: F (19, 76) = 2.433, P = 0.0034 (2-way repeated-measures ANOVA). N.S. P > 0.05, *P < 0.05, **P < 0.01 (Tukey’s multiple comparisons test comparing main effect within treatment groups for each time point). All statistically significant post-hoc comparisons are shown. There was no significant interaction between Time and Treatment.

To address whether SNI altered how the input from *Mrgprd^Cre^* lineage neurons was integrated in the central nervous system, we performed RTPA assays. Once again, at baseline, the amount of time that naïve *Mrgprd^Cre^;* Ai32 mice spent on the blue-light side was not different than that of control mice, confirming that activity in *Mrgprd^Cre^* lineage neurons is not aversive in the absence of injury. In contrast, however, optogenetic activation of *Mrgprd^Cre^* lineage neurons in mice with SNI caused significantly more avoidance than activation of these neurons in naïve mice (Fig. 4D). This finding — in essence, optogenetic allodynia — suggested that activity in non-peptidergic afferents becomes aversive following nerve injury. Moreover, the lack of sensitization post-injury in *Mrgprd^Cre^* lineage afferents to optogenetic stimulation implied that central mechanisms must mediate optogenetic allodynia due to SNI.

### Modulation of spinoparabrachial neurons by Mrgprd^Cre^ lineage afferent input

To investigate the spinal circuitry underlying this phenomenon, we performed whole-cell patch-clamp recordings of lamina I neurons using the *ex vivo* somatosensory preparation (Fig. 5A). Optogenetic stimulation of either the peripheral or central terminals gave rise to EPSCs onto recorded neurons in lamina I, albeit with different latencies (Fig. 5B). We also noted that the EPSC magnitude was larger when the optogenetic stimulation occurred at the skin rather than at the dorsal horn (Fig. 5C). However, there were several instances in which optogenetic stimulation at the dorsal horn was efficacious, whereas stimulation at the skin was not, likely reflecting the fact that not all recorded neurons had receptive fields in the skin of the dorsal hindpaw. For this reason, we selected central stimulation for subsequent experiments.

**Figure 5.**
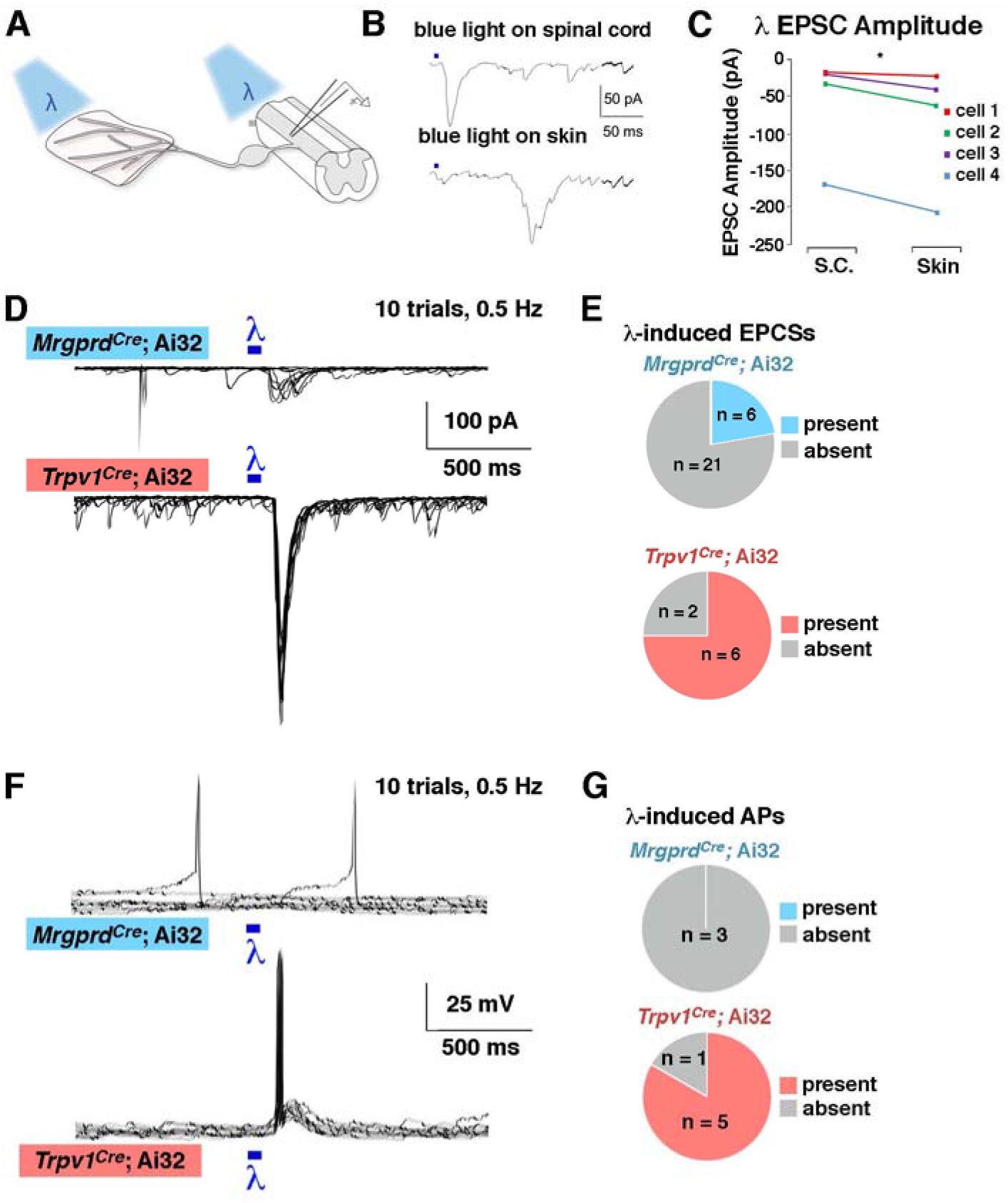
*Lamina I neurons receive strong input from Trpv1^Cre^ lineage afferents but weak input from Mrgprd^Cre^ lineage afferents* (A) Schematic of recordings with optogenetic stimulation at either the peripheral or central terminals. (B - C) Representative traces and quantification of EPSC amplitude from lamina I neurons following optogenetic stimulation at either the skin or the spinal cord. * indicates p < 0.05 (paired Student’s t-test). (D – E) Representative voltage clamp recordings (V_H_ = - 70) and quantification of the proportion of lamina I neurons that show EPSCs upon optogenetic stimulation at the spinal cord in *Mrgprd^Cre^;* Ai32 and *Trpv1^Cre^;* Ai32 mice. (F – G) Representative current clamp recordings and quantification of the proportion of lamina I neurons that show action potentials upon optogenetic stimulation at the spinal cord in *Mrgprd^Cre^;* Ai32 and *Trpv1^Cre^;* Ai32 mice. For D and F, the responses to ten stimulus presentations are superimposed.

We found that optogenetic activation of the central terminals of *Mrgprd^Cre^* lineage neurons gave rise to EPSCs in only 6 of 27 random lamina I neurons (Figs. 5D-E). In contrast, optogenetic activation of the central terminals of *Trpv1^Cre^* lineage neurons gave rise to EPSCs in 6 of 8 random lamina I neurons, which represented a significantly higher proportion relative to *Mrgprd^Cre^* (Fisher’s exact test; p < 0.05). Moreover, optogenetic activation of afferents in *Trpv1^Cre^;* Ai32 mice was sufficient to generate action potentials in 5 of 6 neurons that exhibited EPSCs, whereas the optogenetic activation in *Mrgprd^Cre^;* Ai32 mice was insufficient to generate action potentials in lamina I cells (Figs. 5F-G), which again represented a significantly lower proportion than that observed upon activation of *Trpv1^Cre^* lineage neurons (Fisher’s exact test; p < 0.05). Since randomly targeted neurons are predominantly interneurons, these observations suggest that *Mrgprd^Cre^* lineage neurons do not provide strong excitatory input, either directly or indirectly, onto lamina I interneurons.

### Contribution of Mrgprd^Cre^ lineage afferents to aversion following spared nerve injury

Our behavioral experiments suggested that *Mrgprd^Cre^* lineage input only became aversive following SNI. To investigate mechanisms that may contribute to this injury-induced change, we performed whole-cell recordings from lamina I SPB neurons in naïve or SNI animals (Fig. 6A). In naïve mice, less than half of lamina I SPB neurons displayed EPSCs in response to brushing of the skin with a paint brush, and this input led to action potentials in only 2 of 12 cells (Fig. 6B). Following nerve injury, however, 100% (13 of 13) lamina I SPB neurons showed EPSCs in response to the same dynamic, low threshold input. Moreover, brushing of the skin from SNI mice resulted in action potentials in ~70% of LI SPB neurons in SNI mice, representing a 4-fold increase compared to naïve mice. These results raise the possibility that allodynia in SNI mice may be caused by increased LI spinal output to low threshold stimuli, such as brush. Next, we began investigating the contribution of *Mrgprd^Cre^* lineage neurons to this abnormal spinal output. In naïve mice, 7 of 12 lamina I SPB neurons showed optogenetically-induced EPCSs upon stimulation of *Mrgprd^Cre^* lineage afferents at 2 Hz. Following SNI, however, this proportion increased significantly, with 12 of 13 lamina I SPB neurons now responding to optogenetic stimulation (Fig. 6C). It should be noted, however, that this emergent *Mrgprd^Cre^* lineage neuron input was still rarely sufficient, in itself, to elicit an action potential in the recorded output neuron. Thus, although *Mrgprd^Cre^* lineage neurons are unlikely to be the only afferents involved in allodynia, they may nevertheless provide an important contribution. To gain insight into the underlying mechanism, we analyzed the EPCSs in more detail. Although a higher fraction of lamina I neurons showed light-induced EPSCs following SNI, the responses in individual neurons to optogenetic stimulation were similar and we saw no evidence of synaptic sensitization. Instead, we found that the average response latency to optogenetic stimulation was significantly increased following SNI (Fig. 6D). This elevated latency implies that the enhanced number of responders to *Mrgprd^Cre^* lineage stimulation occurs through an emergent polysynaptic pathway that is normally silent in naïve mice (Fig. 6E).

**Figure 6.**
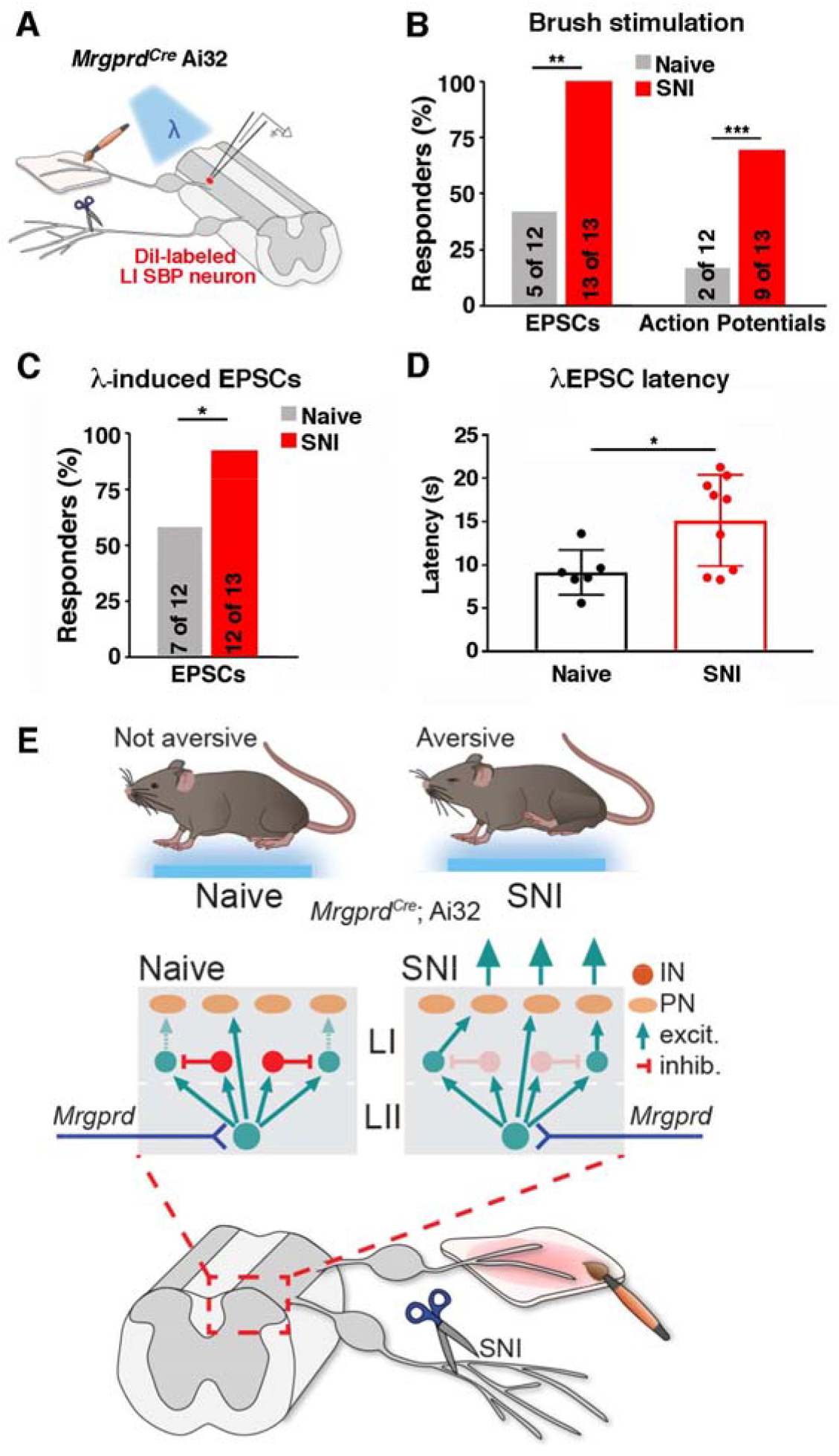
*Increased proportion of lamina I SPB neurons are activated following brush and optogenetic stimulation of Mrgprd^Cre^ lineage afferents neurons following SNI.* (A) Schematic showing patch clamp recordings of lamina I SPB neurons with brush stimulation or optogenetic stimulation. (B) Percent of lamina I SPB neurons that show either EPSCs or action potentials (APs) in response to brushing of the skin in naïve mice or those with SNI. ** indicates P < 0.01 and *** indicates P < 0.001 relative to naïve (one-sided Chi-squared test). (C) Percent of lamina I SPB neurons that show EPSCs upon optogenetic stimulation of *Mrgprd^Cre^* lineage neurons in naïve and SNI mice. * indicates p < 0.05 (one-sided Chi-squared test). (D) EPSC latency to optogenetic stimulation of *Mrgprd^Cre^* lineage neurons in naïve mice or those with SNI. * indicates P < 0.05 relative to naïve (unpaired, Student’s t-test). (E) Proposed model of spinal circuitry activated by *Mrgprd^Cre^* lineage neurons before and after SNI.

## DISCUSSION

Our study provides evidence that activation of *Mrgprd^Cre^* lineage afferents in a naïve state is not aversive, but becomes aversive in the context of chronic neuropathic pain caused by SNI. At the circuit level, we found that lamina I SPB neurons show significantly more brush-induced activity following nerve injury, which is an attractive mechanism to account for the phenomenon of mechanical allodynia. Moreover, *Mrgprd^Cre^* lineage afferents may contribute to this abnormal spinal output after SNI because more lamina I SPB neurons receive *Mrgprd^Cre^* lineage afferent input following optogenetic activation relative to naïve controls. Finally, this increased input appears to be due to an emergent polysynaptic pathway, as evidenced by an increase in the average response latency to optogenetic stimulation of *Mrgprd^Cre^* lineage afferents. Altogether, our study suggests that SNI gives rise to changes in spinal circuitry that enables increases brush-evoked activity in lamina I SPB neurons, and that this effect is likely due, at least in part, to increased input from *Mrgprd^Cre^* lineage afferents through a polysynaptic neural pathway that is normally silent in naïve mice.

Previous studies have identified many mechanisms of injury-induced plasticity in the dorsal spinal cord [2; 28]. Our data suggest that one of these mechanisms involves the engagement of a polysynaptic circuit that emerges following SNI, as evidenced by the increased average latency of input from non-peptidergic afferents. These findings are consistent with the idea that allodynia following nerve injury is caused by disinhibition, which results in low threshold stimuli abnormally reaching nociceptive pathways in the superficial dorsal horn [25].

The results presented here show that low frequency activation of cutaneous *Mrgprd^Cre^* lineage afferents does not induce aversive behaviors. This lineage represents the majority of C-fibers innervating the skin. Thus, an important question is, what information is being conveyed during low frequency activation of these fibers? One potential answer to this question is found in a recent study looking at parallel ascending SPB pathways projecting to distinct subsets of lateral parabrachial sub-nuclei [7]. They found that activation of one of these SPB pathways (GPR83, which is associated with cutaneous mechanosensation) could have either a positive or negative valence depending on the intensity of the optogenetic stimulation. Therefore, low intensity activation of these fibers could possibly evoke a sensation of innocuous touch.

One of the limitations of this type of study is that optogenetic stimulation does not perfectly mimic natural stimulation. In this case, we found that light-mediated activation gave rise to action potentials in *Mrgprd^Cre^* lineage afferents that was similar in frequency to that observed upon stimulation with low threshold stimuli (e.g., 10 mN), but we did not achieve the instantaneous firing frequencies that are typically observed upon stimulation with high threshold stimuli (e.g., 50 mN). For this reason, we cannot exclude the possibility that increasing the frequency of action potentials generated by optogenetic stimulation would alter its valence. Nevertheless, activation of this population at low frequency (2 Hz) is clearly not aversive in naïve mice, whereas activation of afferents that includes the cutaneous peptidergic C-fibers is. This important distinction raises the possibility that the bona fide nociceptors—those whose main function is to warn the organism of tissue damage—are primarily represented by peptidergic C-fibers, and that the function of non-peptidergic C-fibers is likely to be much more nuanced and contextdependent.

An unexpected finding of our work was that control mice lacking ChR2 expression nevertheless showed a modest degree of avoidance for the blue-light side of the RTPA apparatus that emerged after repeated exposure. Since both sides of the chamber produce equally low levels of thermal radiation, it is likely that this avoidance was in response to visual, rather than cutaneous, input. Moreover, mice avoided blue over amber light despite similar lux (illuminance), suggesting that it is the wavelength of light that contributes to the avoidance. We found that the avoidance observed in control mice was most pronounced when light was on continuously. In contrast, the optogenetically-induced aversion in *Trpv1^Cre^;* Ai32 mice was most pronounced when the light was on transiently, flashing at 2 Hz. Because optogenetic stimulation in the wind-up mode gave rise to the highest degree of aversion and thus seemed to be the most effective, we selected this stimulation paradigm for the behavioral experiments involving SNI. However, we note that it will be important to investigate a wide range of stimulation frequencies, including ones that we were not able to achieve with the current tools, to get a more complete picture of the relationship between stimulation frequency and perceptual percept.

Despite the caveat that mice are able to see cutaneous optogenetic stimulation through the floor in RTPA studies, we feel that this approach is still preferable to the implantation of fibers or LEDs through a surgical manipulation because the activation of cutaneous afferents with light does not involve tissue damage, which is likely to be confounding in pain studies. In this regard, it is noteworthy that sham-operated mice that expressed ChR2 in *Mrgprd^Cre^* lineage afferents also showed a trend toward avoidance of the blueside of the RTPA chamber that was greater than that observed in naive mice, though not as pronounced as that observed in SNI mice. This intermediate phenotype of sham treatment suggests that the surgical procedure involving the cutting of skin and muscle is in itself an injury model that may be sufficient for ongoing changes in spinal circuitry.

Just as reported previously [16], we found that Mrgpra3-expressing afferents and Sst-expressing afferents are derived from the Mrgprd lineage. Intriguingly, all three of these afferent subtypes show predominant (though not exclusive) cutaneous targeting, and all three have been implicated in itch [11]}[15; 19; 24]. We did not observe itch behaviors — scratching or biting — upon optogenetic activation of *Mrgprd^Cre^* lineage afferents in our study. Curiously, significant itch behavior has only been reported upon chemogenetic (and not optogenetic) activation of these populations [11; 19; 22]. We speculate that it is not simply the type of afferent input, but also the pattern and the frequency of activation that is interpreted by the nervous system to differentially represent itch, pain, and some aspects of touch. Understanding this integration is of fundamental importance to our basic understanding of somatosensation and may one day lead to the development of improved strategies for the treatment of pain.

## ACKNOWLEDGMENTS

The research reported in this publication was supported by the National Institute of Neurological Disorder and Stroke of the National Institutes of Health under Award Number R01 NS096705 to H.R. Koerber and the National Institute of Arthritis and Musculoskeletal and Skin Diseases of the National Institutes of Health under Award Number R01AR063772 to S.E. Ross.

